# Green, orange, red, and far-red optogenetic tools derived from cyanobacteriochromes

**DOI:** 10.1101/769422

**Authors:** Jaewan Jang, Sherin McDonald, Maruti Uppalapati, G. Andrew Woolley

**Affiliations:** Department of Chemistry, University of Toronto, 80 St. George St., Toronto, ON, M5S 3H6, Canada; Department of Pathology and Laboratory Medicine, University of Saskatchewan, Saskatoon, Saskatchewan S7N 5E5, Canada

**Keywords:** photo-control, GAF, optogenetics, CBCR, phage display

## Abstract

Existing optogenetic tools for controlling protein-protein interactions are available in a limited number of wavelengths thereby limiting opportunities for multiplexing. The cyanobacteriochrome (CBCR) family of photoreceptors responds to an extraordinary range of colors, but light-dependent binding partners for CBCR domains are not currently known. We used a phage-display based approach to develop small (~50-residue) monomeric binders selective for the green absorbing state (Pg), or for the red absorbing state (Pr) of the CBCR *Am1_c0023g2* with a phycocyanobilin chromophore and also for the far-red absorbing state (Pfr) of *Am1_c0023g2* with a biliverdin chromophore. These bind in a 1:1 mole ratio with K_D_s for the target state from 0.2 to 2 μM and selectivities from 10 to 500-fold. We demonstrate green, orange, red, and far-red light-dependent control of protein-protein interactions *in vitro* and also *in vivo* where these multicolor optogenetic tools are used to control transcription in yeast.

## Introduction

Optogenetic tools that enable the control of protein-protein interactions are highly desirable for numerous cell biology applications (1, 2). Most available tools of this kind are naturally occurring photoswitchable binding partners (*e.g.* Cry2/CIBN, PhyB/PIF3, BPhP1/PpsR2) (3–5). Protein engineering approaches have been employed to develop improved photoswitchable protein-protein interaction modules based primarily on LOV domains (1, 6, 7). While powerful, current tools have numerous limitations. For example, some of them (*e.g.* Gigantea, Cry, Phy) (8) are large and/or oligomeric and thus may perturb the folding and function of the proteins they are fused to (3, 9). While there are numerous examples of light-induced protein-protein association, light induced dissociation is rare (AsLOV/zDark and cPYP/BoPD (10, 11)). Most significantly, however, the currently available range of colors is incomplete - the large phytochrome systems respond to red/far red-light (3) but the vast majority of optogenetic tools for controlling protein-protein interactions respond to blue light (www.optobase.org)(12). As a result, opportunities for multiplexing optogenetic tools, or for using optogenetic tools together with the wide range of available fluorescent reporters are limited (3).

The cyanobacteriochromes (CBCRs), a superfamily of photoreceptor proteins discovered about ten years ago in cyanobacteria (13, 14) respond to a wide range of colors. These proteins typically consist of multiple subdomains called GAF domains (cGMP-phosphodiesterase/adenylate cyclase/FhlA) linked in tandem (15). A conserved cysteine in the GAF domain covalently binds a bilin chromophore (typically phycocyanobilin (PCB)) via a thioether bond to the C3^1^ atom and irradiation triggers E/Z isomerization across the C15 double bond of the chromophore (16). Changes in the GAF protein environment around the bilin chromophore tune the wavelength of absorption to an extraordinary degree (14, 16). Recent discoveries have identified members of the CBCR family that respond to virtually any wavelength in the visible range from the far-red to the UV (14, 17–21). In addition to their wide range of colors, GAF domains are small (<200 residues) and can function as monomers (22, 23) – both desirable features for optogenetic tools.

Photoisomerization of the bilin chromophore leads to conformational changes in the GAF domain (24). A recent structural study of the red/green cyanobacteriochrome, NpR6012g4, showed that upon isomerization, a small segment of the protein near the chromophore changes from an irregular to a helical structure and subtler structural rearrangements are propagated throughout the protein (25). It has been hypothesized that GAF conformational changes modulate the dimerization propensity of CBCRs and/or the conformation of pre-formed dimers and this, in turn, regulates the activity of C-terminal effector domains (23). Conformational changes propagated via the C-terminal linker have also been proposed to directly alter the conformation of the attached effector domain without stable dimerization (26). No naturally occurring light-dependent binding partners for CBCRs have been reported.

Recently we developed a phage-display approach to finding binding partners for photoswitchable proteins for which no known binding partner exists (10). The binding partners were developed by randomizing a surface on the small stable 3-helix bundle GA domain using a customized codon set (10). Here we apply a slightly modified approach to discover binding partners for a GAF domain from the CBCR family. As a target we chose *Am1_c0023g2*, a red/green GAF domain from *Acaryochloris marina*, that can bind two distinct bilins - PCB or biliverdin (BV) (27). When bound to PCB, *Am1_c0023g2* forms a thermostable red-absorbing (Pr) state. When irradiated with red light (680 nm), the Pr state converts to a green absorbing state (Pg). In the dark the Pg state exhibits very slow thermal reversion (τ½ >24h, 25°C) to the Pr state. Irradiation of the Pg state with green light (525 nm) rapidly and quantitatively converts the protein back to the Pr state. When *Am1_c0023g2* binds the more extensively conjugated BV chromophore, it forms a thermostable far-red absorbing (Pfr) state. The Pfr state can be converted using 750 nm light to an orange absorbing (Po) state, which reverts thermally to the Pfr state (τ½ = 180 min, 25°C) or can be rapidly switched back to the Pfr state with orange light (595 nm). Thus, by changing the nature of the bilin chromophore loaded onto the protein, *Am1_c0023g2* can be made responsive to either green, orange, red or far-red light. Optogenetic tools that respond to green light are currently limited to complex cobalamin based systems (28, 29) and CcaS/CcaR or cPAC systems that have predefined functions - gene expression (30), or adenylyl cyclase activity respectively (26). In addition, small, monomeric optogenetic tools that operate with far-red light are useful since far-red absorption provides deep tissue penetration ability for use in living animals (31). Proteins that operate using BV are of particular interest for studies in living animals where BV, though not PCB, is available via heme metabolism (27, 32).

## Materials and Methods

### Production of Am1_c0023g2 PCB and BV

For production of *Am1_c0023g2 in vitro* the gene was subcloned into a pET24b vector containing a C-terminal poly His (6x) tag and apo-protein was expressed in *E. coli* BL21(DE3). A two-fold molar excess of purified PCB or BV (Frontier Scientific) in DMSO was added as described in the SI. For *in vivo* chromophore reconstitution, the gene was subcloned into a pBAD-HisC vector with a C-terminal 6x His-tag. *E. coli* strain LMG194 containing pPL-PCB (a kind *gift* from J.C. Lagarias) or pWA23h (a kind gift from V.V. Verkhusha) was transformed with pBAD-*Am1_c0023g2* and the protein was expressed and purified as described in the SI.

### Phage display-based screening

A M13 phage pVIII library based on the GA domain described previously (10) was used to find binders for the Pg, Pr, Po or Pfr states of *Am1_c0023g2* using the detailed protocol described in the SI. The GA domain phage library was first depleted for binders to the apo protein. A second negative selection was carried out against the off-target state, followed by a positive selection on target. The resulting library of positive clones obtained after 3 rounds of selection was subcloned into a pIII phagemid and ~100 single clonal phage from each of the positive pools were screened for binding using phage-based ELISAs (33). For affinity maturation, biased libraries were constructed in the pIII display format, based on doped oligonucleotides specific for lead clones BAm-green1.0 and BAm-red1.0 as described in the SI.

### In vitro characterization of binders

Selected binders were subcloned from pIII phagemids into the pET24b expression vector containing a C-terminal poly His (6x) tag and expressed and purified from *E. coli*. Size exclusion binding assays were performed by mixing *Am1_c0023g2* PCB or BV (80 μM) with each binder (80 μM) in PBS and injecting onto a Superdex 75 10/300 GL size exclusion column maintained in the dark or while irradiating with either red (680 nm), green (525 nm), or far red (750 nm) light. ITC experiments were performed using a MicroCal VP-ITC MicroCalorimeter. Titration experiments were performed at 25°C. The syringe contained the binder, BAM-red1.0 or BAm-green1.3, at 550 μM or 750 μM, respectively. The cell contained either *Am1_c0023g2* PCB exo (45 μM), *Am1_c0023g2* PCB endo (42 μM), *Am1_c0023g2* BV exo (45 μM) or *Am1_c0023g2* BV endo (42 μM). Thermogram data were integrated using NITPIC (34) and binding analysis was carried out using SEDPHAT and recommend protocols (35, 36).

### Yeast two-hybrid assays

The strains Y2HGOLD and Y187 were purchased from Clontech. pGAL4AD-x and pGAL4BD-y plasmids were purchased from Addgene (28246 and 28244) as pGAL4AD-CIB1 and pGAL4BD-Cry2 (4). The *Am1_c0023g2* gene was PCR amplified and inserted (Gibson Assembly) into either the pGAL4AD or the pGAL4BD vector by replacing a CIB1 or Cry2 gene, respectively. Binder constructs were also subcloned into both plasmids using the same protocol. Strains containing pGAL4AD-x and pGAL4BD-y, respectively, were mated and selected on synthetic media lacking leucine and tryptophan. A single colony was picked and used to inoculate media containing 10 μM PCB or 40 μM BV (Frontier Scientific). Cultures were grown under different illumination conditions (green, red, far-red, dark), cells were harvested and β-galactosidase activity was measured following the manufacturers protocol as described in the SI. The experiment was performed in quadruplicate.

## Results and Discussion

### Am1_c0023g2 engineering, expression and reconstitution

Cyanobacteriochromes occur naturally as multi-domain proteins that may dimerize to varying extents (23). Since use as an optogenetic tool for controlling protein-protein interactions is simpler if the interacting protein partners are monomers, we sought to prevent dimerization of *Am1_c0023g2*. Dimerization of cyanobacterial GAF domains is analogous to that of PHY-GAF-PAS domains from bacterial phytochromes (24). Previous work successfully monomerized a bacterial phytochrome by preventing a coiled-coiled interaction involving the C-terminal helix (32). We therefore mutated Leu159 on the C-terminal helix to Lys and also truncated the N-terminal helix of *Am1_c0023g2* to prevent homodimer formation (see sequences in SI Fig. S1).

We expressed this variant of *Am1_c0023g2* with a C-terminal His-tag using an *E. coli* expression system, either with exogenous addition of PCB or BV or via coexpression of the enzymes to make each bilin (38). The protein eluted at a size corresponding to a monomeric, 20 kDa protein when analyzed using size exclusion chromatography (SEC) (Fig. S2). Spectra obtained matched those described by Narikawa and colleagues (27) except that the maximum wavelength of the Pg state observed via exogenous addition of PCB was red-shifted compared to that obtained for the Pg state of *Am1_c0023g2* reconstituted via coexpression of heme oxygenase (HO1) and phycocyanobilin:ferredoxin oxidoreductase (PcyA) to produce PCB *in vivo* (Fig. 1, Fig. S3). Spectra for BV-loaded *Am1_c0023g2* were insensitive to whether the bilin was added exogenously or endogenously (Fig. 1, Fig. S3). Differences in chromophore absorption spectra that depend on the mode of chromophore loading have been noted previously for other CBCRs (39, 40) and have been attributed to local conformational differences in the bilin binding pocket that may occur depending on whether chromophorylation occurs co-translationally or post-translationally (39). Presumably the same sensitivity to local environment that leads to the extraordinary degree of color tuning seen with the CBCR photoreceptor family can also lead to spectral differences if conformational changes in the bilin binding pocket occur with protein expression changes (*e.g.* changes in host organism, rate of synthesis, temperature etc.). Additionally, a consistent observation for both exogenously-assembled and endogenously-assembled preparations is that significant amounts of apoprotein are often present (18, 40). These features of the CBCR family are important to bear in mind when employing CBCRs as optogenetic tools in heterologous systems since the mode of chromophore loading (*e.g.* adding to cells in culture (41), or via coexpression of chromophore biosynthesis genes (40, 42) may lead to a mixture of apoprotein and distinct subsets of holoproteins in specific cases. Since we anticipate that initial applications of multicolour GAF-based optogenetic tools will be cells in culture where exogenous addition of chromophore is straightforward and provides a convenient way of commencing an experiment (10, 41) and because the exogenous reconstitution of *Am1_c0023g2* with PCB and BV reliably produced ~100% holo protein in our hands, we first focused on finding binding partners to the exogenously produced forms of the proteins. In addition, we aimed to find binding partners that would not recognize the apoprotein.

**Figure 1.**
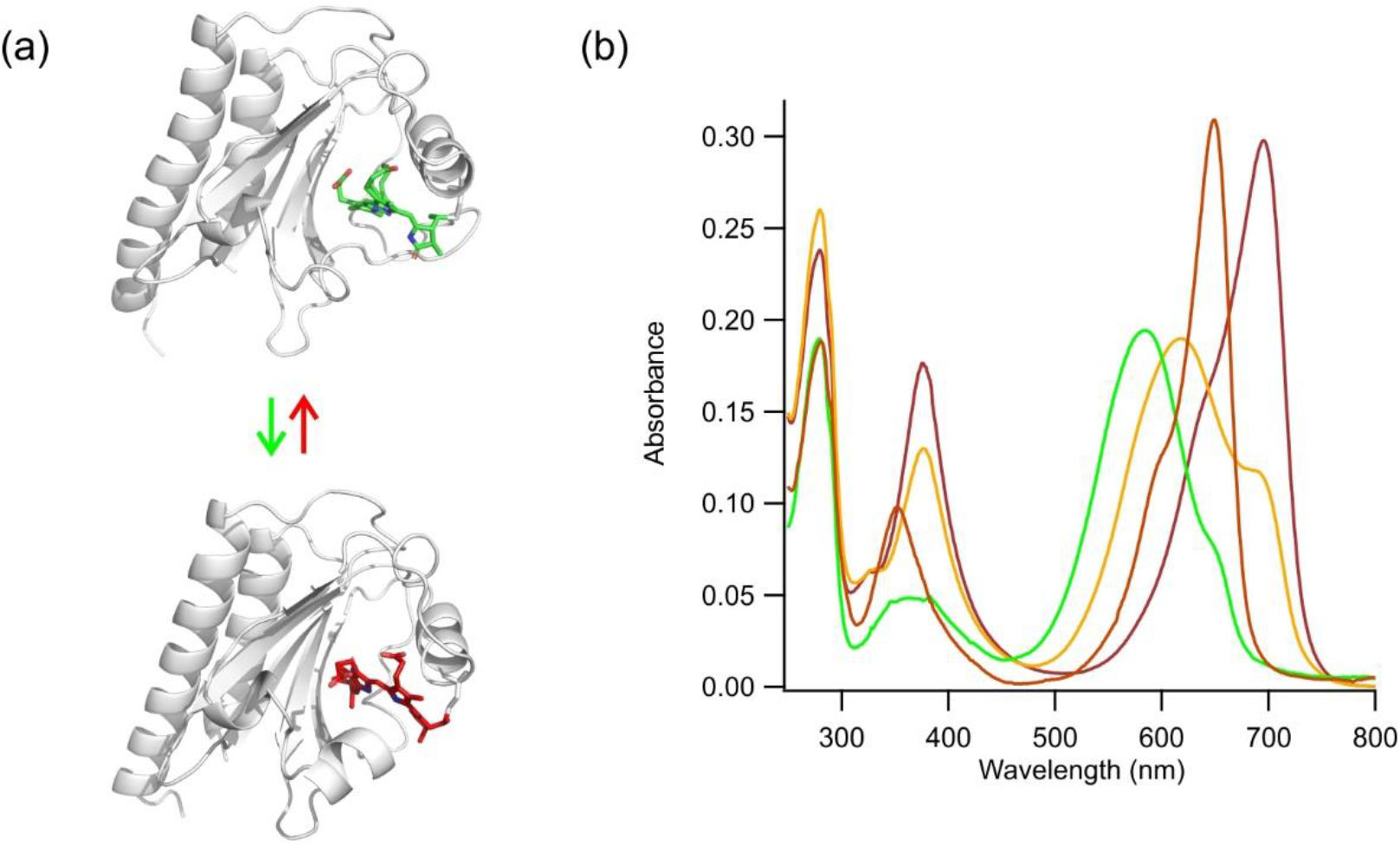
(a) Modeled structures of Pg and Pr states of *Am1_c0023g2* PCB. Structures were generated by threading the *Am1_c0023g2* sequence through models of NpR6012g4 (PDBIDs: 6BHN Pr state; 6BHO, Pg state (25)) using Phyre2 (37) (b) Absorbance spectra of *Am1_c0023g2* with exogenous PCB incorporation in the Pr state (red line) and Pg state (green line) and with exogenous BV incorporation in the Pfr state (far-red line) and the Po state (orange line).

### Phage-display based discovery of binding partners for Am1_c0023g2

In previous work we successfully used pVIII displayed libraries of the 3-helix bundle GA domain to find binders for both the blue-light and dark-adapted states of LOV and PYP photoreceptors (10). We used the same library to search for binders to distinct conformational states of apo- and holo- *Am1_c0023g2* with either PCB or BV chromophores. First, we carried out a negative selection step to deplete the phage pool for binders to the apo-protein. To select for binders to the Pg state, a second negative selection step to remove binders for the Pr state was carried out by exposing the library to a plate containing holo-*Am1_c0023g2* PCB in the Pr state. Unbound phage from this plate were then transferred to a third plate with holo-*Am1_c0023g2* PCB in the Pg state. This plate was washed extensively, then binders were eluted and amplified. This three-step selection process was repeated two more times. Selection for binders to the Pr, Po, and Pfr states was carried out in an exactly analogous manner. The resulting phage pools were then subcloned into a pIII display phagemid as described previously (10). Since there are fewer copies of the GA domain displayed in the pIII fusion compared to the pVIII fusion context per phage this step may select for higher affinity binders by decreasing the likelihood of avidity-mediated binding. In addition, clones that function in both pIII and pVIII contexts are expected to be less sensitive to the presence of surrounding phage proteins for their function. Perhaps because of the subtle conformational change between photo-states that was being selected for, very limited enrichment of phage libraries was observed at the pool level. We therefore screened 96 single clones using phage ELISA assays to search for photo-state selective binders.

Clones were selected based on their apparent affinity and selectivity in phage-based ELISA assays (Fig. 2). The best clone for state selective binding to *Am1_c0023g2* PCB in the Pg state, designated BAm-green1.0 (for binder of *Am1_c0023g2-*Pg) showed a 13-fold better binding to the Pg versus the Pr state. The best clone for selective binding to holo-*Am1_c0023g2* PCB in the Pr state, designated BAm-red1.0 (for binder of *Am1_c0023g2-*Pr) showed 7-fold better binding for Pr over Pg. Both clones showed no detectable binding to the apo form of *Am1_c0023g2* (Fig. 2). While no Po-state selective binders *Am1_c0023g2* BV were found, several sequences were found to be selective for the Pfr state (Fig. 2c). Interestingly, the Pr-state binder BAm-red1.0 was also found to bind the Pfr state with similar affinity and selectivity to those binders directly selected for the Pfr state. In contrast, BAm-green1.0 did not bind the Po state of GAF2-BV (Fig. 2c). We therefore opted to pursue BAm-red1.0 as both a Pr and a Pfr state binder.

**Figure 2.**
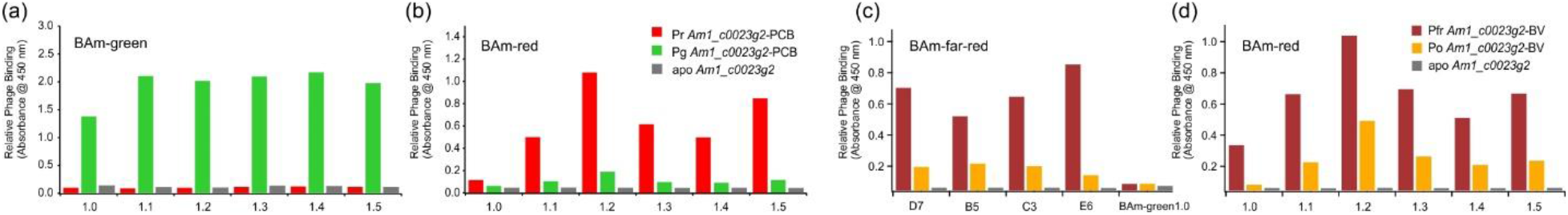
Phage-based ELISAs of BAm-green, BAm-red, and BAm-far-red sequences and their affinity matured variants. (a) BAm-green 1.0 was selected against the Pg state of *Am1_c0023g2-*PCB. BAm-green1.1-1.5 are affinity-matured variants. (b) BAm-red 1.0 was selected against the Pr state of *Am1_c0023g2* PCB. BAm-red1.1-1.5 are affinity-matured variants. Although the signal from BAm-red1.0 is lower, the specificity is higher than for 1.1-1.5. (c) BAm-far-red clones were selected against the Pfr state of *Am1_c0023g2* BV. No clones selective for the Po state of *Am1_c0023g2* BV were found. BAm-green1.0 does not bind to either state of *Am1_c0023g2* BV. (d) BAm-red and affinity-matured variants selectively bind the Pfr state of *Am1_c0023g2* BV.

In the absence of structural information, it is not clear which of the previously randomized positions of the GA domain scaffold make contact with the target protein in each state. Therefore, we used a soft-randomization approach to improve the affinity of binders (43). We constructed biased pIII phage libraries based on BAm-green1.0 and BAm-red1.0 sequences. Here each of the nucleotides encoding the randomized positions was substituted with a doped mix such that the native nucleotide in the parent DNA sequence was doped at 70% while the rest of the 3 nucleotides occured at 10% frequency each (see Methods). This allows for approximately 40% bias for the native amino acid in each randomized position, while allowing the occurrence of the other 19 amino acids at lower frequencies. Four rounds of selection were carried out using the same protocol as described for the naïve library, then ~24 clones were tested using phage ELISAs. Figure 2 shows the top five clones obtained for each affinity selection target. With BAm-green, most new clones showed improved affinity with no loss in specificity (Fig. 2). Among the variants, BAm-green-1.3 proved to have the highest affinity with no loss in selectivity. BAm-red1.0 variants (BAm-red-1.1-1.5) also showed improved affinity (Fig. 2), but similar or reduced specificity for binding to the Pr state over the Pg state, or the Pfr state vs. the Po state (Fig. 2). Affinity matured versions of BAm-red1.0. may be suitable for applications that can tolerate some basal activity. Based on these results BAm-green1.3 and BAm-red1.0 (which also binds the Pfr state) were selected for detailed characterization *in vitro*.

### In vitro characterization of Am1_c0023 binding partners by SEC

Binder sequences selected from pIII ELISAs were cloned into expression vectors coding for a C-terminal His-tag, expressed in *E. coli* and purified via Ni-NTA affinity chromatography, followed by size-exclusion chromatography (SEC). Based on retention volumes in SEC, all binders were monomeric (Fig. S4). We then used SEC to detect light-dependent interactions between *Am1_c0023g2* and BAm variants. When BAm-green-1.3 was mixed with *Am1_c0023g2* PCB in the Pr state, two peaks were observed in the chromatogram with elution volumes matching those observed when *Am1_c0023g2* PCB and BAm-green1.3 were injected individually (Fig. 3a). In contrast, when BAm-green1.3 was mixed with *Am1_c0023g2* PCB in the Pg state, a new peak eluted with a smaller elution volume. Analysis of elution fractions using SDS-PAGE confirmed the earlier peak was a complex of *Am1_c0023g2* PCB Pg and BAm-green1.3. Exactly opposite behavior was observed with BAm-red1.0 confirming red-light dependent complex formation with *Am1_c0023g2* PCB in the Pr state.

**Figure 3.**
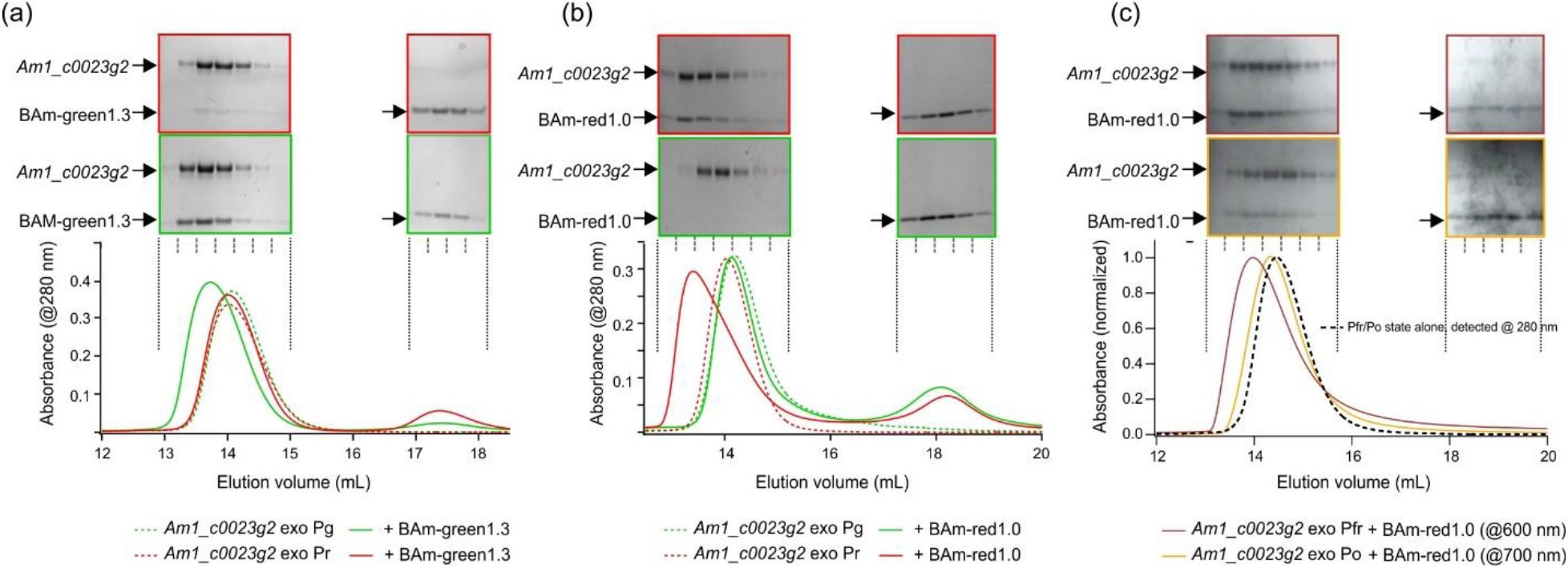
Size exclusion chromatography of *Am1_c0023g2* forming light-dependent complexes with BAm-green1.3 or BAm-red1.0. (a) PCB-loaded *Am1_c0023g2* exo in the Pg state forms a complex with BAm-green1.3 that elutes earlier than *Am1_*c0023g2 alone. SDS-PAGE of column fractions shows this complex contains both proteins. Complex formation is not observed with the Pr state. (b) PCB-loaded *Am1_c0023g2* exo in the Pr state forms a complex with BAm-red1.0 and does not with the Pg state. (c) BV-loaded *Am1_c0023g2* exo in the Pfr state forms a complex with BAm-red1.0. Rapid relaxation from Po to Pfr in the presence of BAm-red1.0 leads to some complex formation when Po is mixed with BAm-red1.0 as detected by SDS-PAGE (and absorbance at 280 nm (Fig. S5)). Selective detection of the Po state by measuring absorbance at 600 nm, however, indicates very little shift in elution volume in the presence of Bam-red1.0 as compared to Po alone.

When *Am1_c0023g2* BV in the Pfr state was mixed with BAm-red1.0 complex formation was again observed (Fig. 3c). When *Am1_c0023g2* BV in the Po state was mixed with BAm-red1.0, a significantly smaller shift in elution volume was observed (Fig. 3c). While this result may indicate some affinity of BAm-red1.0 for the Po state, it may also reflect binding to residual Pfr state. As noted above, while thermal relaxation of *Am1_c0023g2* PCB from the Pg to the Pr state is very slow (>24h at 25°C), thermal relaxation of *Am1_c0023g2* BV from the Po state to the Pfr state is significantly faster (τ½ = 180 min at 25°C). Interestingly, we observed that adding BAm-red1.0 enhanced thermal relaxation in a concentration dependent manner for both *Am1_c0023g2* PCB and for *Am1_c0023g2* BV (Fig. S6). Thermal relaxation, like photoisomerization, of these proteins is likely to involve several intermediate states and associated transition states (44). The observation that BAm-red1.0 catalyzes thermal relaxation implies that it binds the Pg state or an intermediate state and stabilizes a rate-determining transition state involved in the thermal back reaction.

We also tested light-dependent interaction of *Am1_c0023g2* that had been reconstituted via coexpression of the enzymes required to make either the PCB or BV chromophore (*Am1_c0023g2* PCB/BV endo). Size-exclusion chromatography of *Am1_c0023g2* PCB endo in the Pg state mixed with BAm-green1.3 showed decreased complex formation compared to that seen with *Am1_c0023g2* PCB exo in the Pg state (Fig. S7). This result indicates that BAm-green1.3 binding is sensitive to the structural change that underlies the shift in the λ_max_ of the *Am1_c0023g2* PCB Pg state observed with exogenously versus endogenously reconstituted protein. In contrast, when *Am1_c0023g2* PCB endo in the Pr state was mixed with BAm-red1.0, complex formation was observed by SEC that was indistinguishable from that seen with *Am1_c0023g2* PCB exo.

### Affinity and stoichiometry of Am1_c0023 – BAm interactions

To quantify the affinity of the binder proteins for their targets, we carried out isothermal titration calorimetry (ITC) measurements. Titration of BAm-green1.3 into an ITC cell containing *Am1_c0023g2* PCB exo in the Pg state produced the thermogram shown in Figure 4a. These data fit well to a 1:1 binding model with a K_D_ of 0.25 ± 0.1 μM and an exothermic heat of binding of 8.2 ± 0.9 kcal/mol. Titration of BAm-green1.3 into a cell containing *Am1_c0023g2* PCB exo in the Pr state produced the thermogram shown in Figure 4b. As described in the methods section, a fraction of the Pr protein is likely switched to the Pg state during loading of the ITC cell and the thermogram observed is consistent with the presence of 10 ± 5% Pg state. The remaining 90% Pr state does not produce a measurable heat of binding with BAm-green1.3. The observed lack of a shift in the SEC elution volume (Fig. 3a) indicates the affinity of BAm-green1.3 for this state must be >100 μM. Thus, BAm-green1.3 exhibits >500-fold selectivity (change in K_D_) for the Pg state over the Pr state of *Am1_c0023g2* PCB exo.

**Figure 4.**
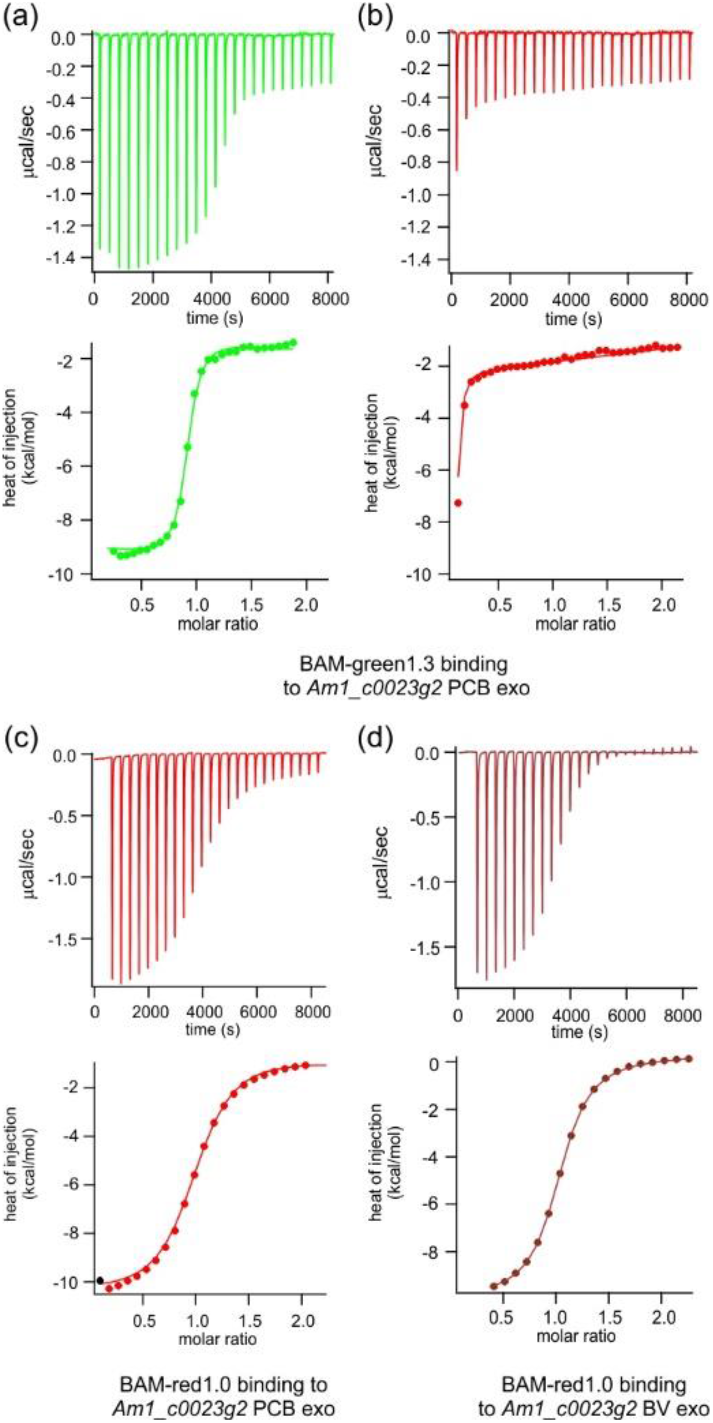
Isothermal titration calorimetry of BAm-green1.3 and BAm-red1.0 binding to *Am1_c0023g2* in a state selective manner. Thermograms are shown in the upper panels and calcuated binding isotherms in the lower panels. (a) BAm-green1.3 (750 μM in the syringe) was titrated into a solution of *Am1_c0023g2* PCB exo (45 μM) in the Pg state and thermogram data fitted to a 1:1 binding model to give K_D_ = 0.25 μM (0.2 - 0.4 μM) and ΔH = −8.2 ± 0.9 kcal/mol. (b) BAm-green1.3 was titrated into a solution of *Am1_c0023g2* PCB exo (45 μM) in the Pr state and thermogram data fitted to a model in which 10% of the Pg state remains and the Pr state is inactive. (c) BAm-red1.0 (550 μM in the syringe) was titrated into a solution of *Am1_c0023g2* PCB exo (45 μM) in the Pr state and thermogram data fitted to a 1:1 binding model to give K_D_ = 1.8 μM (1.4 – 1.9 μM) and ΔH = −10 ± 0.3 kcal/mol. (d) BAm-red1.0 (550 μM in the syringe) was titrated into a solution of *Am1_c0023g2* BV exo (39 μM) in the Pfr state and thermogram data fitted to a 1:1 binding model to give K_D_ = 0.9 μM (0.8 - 1 μM) and ΔH = −10 ± 0.3 kcal/mol.

We then measured binding of BAm-red1.0 to Pr-adapted *Am1_c0023g2* PCB that had been reconstituted either exogenously or endogenously. The thermogram for BAm-red1.0 binding to *Am1_c0023g2* PCB exo is shown in Figure 4b; that for binding to *Am1_c0023g2* PCB endo is shown in Figure S8. As expected, based on the SEC results (Fig. 3) both forms bound similarly. In both cases, data fit well to a 1:1 binding model with a K_D_ of 1.8 ± 0.5 μM (exo) and 0.8 ± 0.5 μM (endo) and with an exothermic heat of binding of 9.5 ± 0.5 kcal/mol. We found that addition of BAm-red1.0 to *Am1_c0023g2* PCB exo in the Pr state caused quenching of Pr-state fluorescence at λ_max_ ~ 670 nm. This effect was used to independently assess the binding affinity of BAm-red1.0 for the Pr state of *Am1_c0023g2* PCB exo (Fig. S9). These data gave a K_D_ of 2.5 ± 1 μM, like that measured by ITC.

ITC analysis of BAm-red1.0 binding to the Pg state of *Am1_c0023g2* PCB was complicated by the fact that the binder enhances thermal relaxation of *Am1_c0023g2* PCB Pg to the Pr state (Fig. S6). The process is slow enough that ITC measurements of BAm-red1.0 binding to Pr state and Pg state samples are qualitatively very different (Fig. S8) but a quantitative measure of the Pg state K_D_ by ITC is difficult. The observed lack of a shift in the SEC elution volume (Fig. 3b) indicates the affinity of BAm-red1.0 for the Pg state must be >100 μM. The observed effect on the thermal relaxation rate (Fig. S6) suggests an affinity of BAm-red1.0 for the Pg state, or an intermediate, of 280 ± 80 μM. Thus, it appears that BAm-red1.0 exhibits a ~200-fold selectivity (change in K_D_) for the Pr state over the Pg state of *Am1_c0023g2* PCB.

Finally, the binding of BAm-red1.0 to the Pfr state of *Am1_c0023g2* BV exo and endo was examined by ITC. These data are shown in Figure 4d and Figure S10. The thermogram data fit well to a 1:1 binding model with a K_D_ of 0.9 ± 0.1 μM (exo) and 0.75 ± 0.5 μM (endo) with an exothermic heat of binding of 10 ± 0.5 kcal/mol. Due to rapid thermal reversion of the Po state to the Pfr state, we could not measure a K_D_ for BAm-red1.0 binding to the Po state.

### Heterologous expression in living cells as optogenetic tools

Having established that BAm-green1.3 and BAm-red1.0 showed selective binding to distinct photo-states of the *Am1_c0023g2* GAF domain *in vitro*, we wished to test whether these light dependent binding partners could function in live cells as optogenetic tools. The yeast two-hybrid system is a well-established protein-protein interaction validation assay and has been used frequently to test naturally occurring as well as engineered optogenetic tools (1, 45). We fused *Am1_c0023g2* to either the GAL4 activating domain (GAL4AD) or the GAL4 DNA binding domain (GAL4BD) and the binding partner to the other (Fig. 5a). If the protein partners associate, transcription of a GAL4-regulated *lacZ* (producing β-galactosidase) reporter should result.

**Figure 5.**
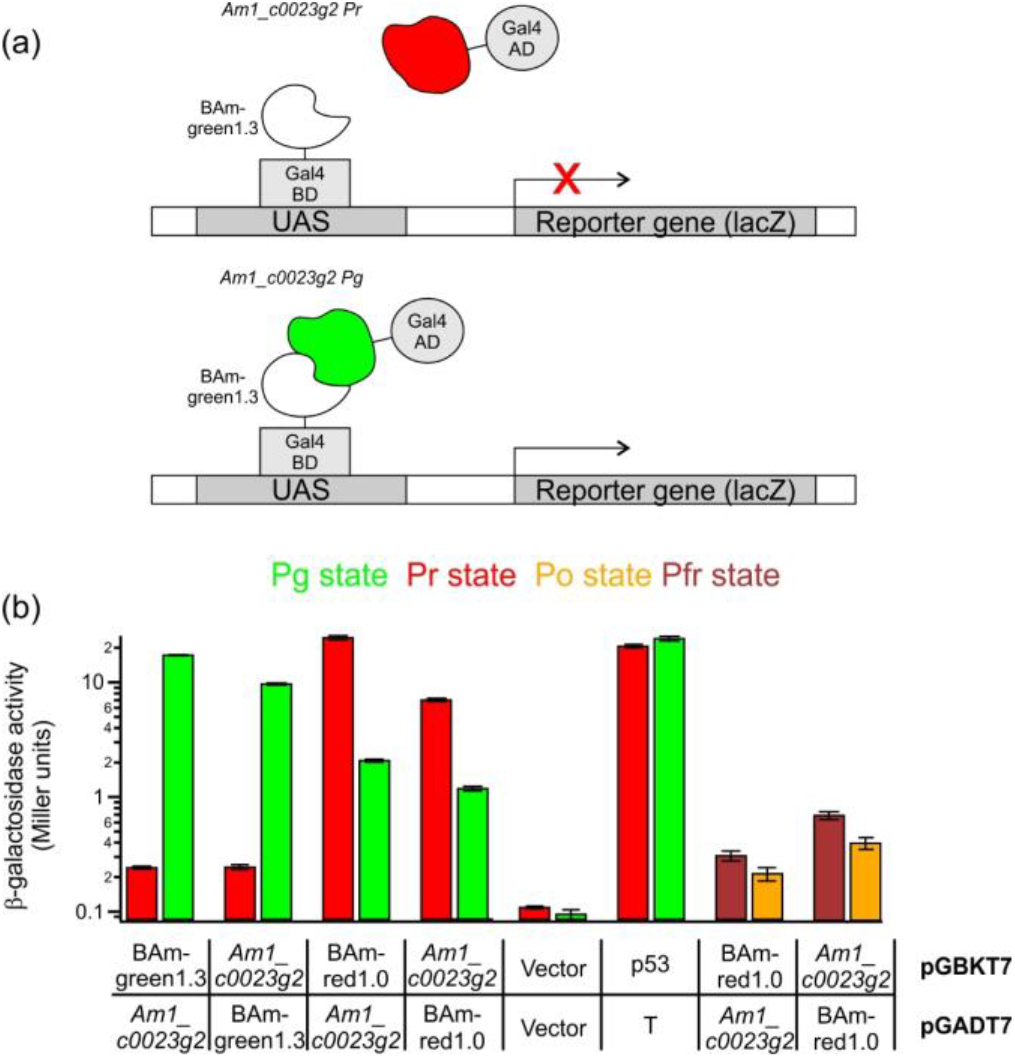
Light induced transcription of a β-galactosidase reporter in yeast. (a) Schematic of yeast two hybrid with GAL4AD and GAL4BD fusions. (b) Light dependent production of β-galactosidase activity in yeast (note log scale). *Am1_c0023g2* PCB or BV and binder constructs were tested for interaction under different illumination conditions to produce either Pg, Pr (PCB) or Po, Pfr states (see methods). Negative control for no interaction (vector only) and positive control (P53/T) are also shown.

Figure 5 shows β-galactosidase activity measured in yeast cells expressing different combinations of the BAm binders and *Am1_c0023g2* in which PCB was added exogenously to the yeast culture. When *Am1_c0023g2* was fused to the GAL4 DNA binding domain and BAm-green1.3 was fused to the GAL4 activating domain, β-galactosidase activity 40% of that of the positive control was observed under red light with <1% activity under green light (a >40-fold difference). When *Am1_c0023g2* was fused to the GAL4 activating domain and BAm-green1.3 was fused to the GAL4 DNA binding domain, β-galactosidase activity ~80% of that of the positive control was observed under red light with still <1% activity under green light (a >70-fold difference).

When the BAm-green1.3 fusions were replaced by BAm-red1.0 fusions the responses to red and green light were reversed as expected. When *Am1_c0023g2* was fused to the GAL4 DNA binding domain and BAm-red1.0 was fused to the GAL4 activating domain, β-galactosidase activity 30% of that of the positive control was observed under green light with < 5% activity under red light (a 7-fold difference). The fold-difference was improved when the orientation of the binding partners was swapped; i.e. when *Am1_c0023g2* was fused to the GAL4 activating domain and BAm-red1.0 was fused to the GAL4 DNA binding domain, β-galactosidase activity ~100% of that of the positive control was observed under green light with <10% activity under red light (a 12-fold difference). Overall these results are consistent with the *in vitro* selectivity measured for BAm-green1.3 and BAm-red1.0 for the Pr and Pg states of *Am1_c0023g2*, i.e. the BAm-green1.3/ *Am1_c0023g2*-Pg state is the most selective interaction. When *Am1_c0023g2* was expressed together with BAm-red1.0 in yeast and BV was added to the media, lower apparent degrees of protein-protein interaction in the on-state were observed (Fig. 5b), which may reflect the lower solubility of BV combined with lower uptake vs. PCB by yeast cells (46) or inefficient formation of holo-*Am1_c0023g2*-BV protein inside yeast. Nevertheless, a reproducible two-fold effect (Pfr vs. Po) on the transcription of β-galactosidase was observed (Fig. 5b).

## Discussion

We have described a phage-display selection approach with negative selection against apo-protein and an affinity maturation step that enabled the development of the first binder proteins for a cyanobacteriochrome (CBCR) GAF domain. One-to-one light-dependent binding to the target state was characterized *in vitro* with affinities between 200 nM and 2 μM for the target photo-state and 10- to 500-fold lower affinity for the off-target photo-state. These interactions can be used to enable red light turn-on and green light turn-off of a protein-protein interaction (BAm-green1.3/*Am1_c0023g2*), or alternatively green light turn-on of a protein-protein interaction and red-light turn-off (BAm-red1.0/*Am1_c0023g2*). Whereas BAm-green1.3 binds differentially to *Am1_c0023g2*-PCB that has been produced via exogenous addition of PCB versus via co-expression of the enzymes that synthesize PCB, BAm-red1.0 binds to the Pr state whether this has been produced exogenously or endogenously. In both cases, functional responses were demonstrated *in vivo* in yeast when PCB was added to the growing yeast cells enabling red/green light inducible transcription. We show that neither binder interacts with apo protein, which is likely to be present in any heterologous system.

By replacing PCB with BV, orange and far-red absorbing forms of *Am1_c0023g2* were obtained. We found that BAm-red1.0 also binds to the Pfr state of *Am1_c0023g2* BV and far-red light produces dissociation of BAm-red1.0/ *Am1_c0023g2* BV. Orange light can also be used to trigger the interaction of BAm-red1.0 and *Am1_c0023g2* BV. This interaction will also occur via thermal relaxation to the Pfr state in the dark. If delocalized far-red light was used to dissociate BAm-red1.0/*Am1_c0023g2* BV, focused orange light could be used to spatially localize a BAm-red1.0/*Am1_c0023g2* BV interaction. While reconstitution of *Am1_c0023g2* BV in yeast was inefficient, protein engineering with these systems is already leading to variants with enhanced capability for accepting BV (47), thereby extending their suitability for use in diverse hosts.

Together, BAm-green1.3 and BAm-red1.0 thus provide new optogenetic components for controlling protein-protein interactions with green, orange, red, and far red light. Because they are based on the GAF domain of a cyanobacteriochrome and a ~50-residue GA three-helix bundle protein, they are the smallest green, orange, red and far-red optogenetic tools thus far identified. The approach described here is expected to be suitable for the generation of other cyanobacteriochrome GAF-based optogenetic tools. The rich diversity of the cyanobacteriochrome family offers further possibilities for even more diverse multicolor optical control.

## References

1. Hallett RA, Zimmerman SP, Yumerefendi H, Bear JE, & Kuhlman B (2016) Correlating in Vitro and in Vivo Activities of Light-Inducible Dimers: A Cellular Optogenetics Guide. ACS Synth Biol 5(1):53–64.

2. Goglia AG & Toettcher JE (2019) A bright future: optogenetics to dissect the spatiotemporal control of cell behavior. Curr Opin Chem Biol 48:106–113.

3. Redchuk TA, Omelina ES, Chernov KG, & Verkhusha VV (2017) Near-infrared optogenetic pair for protein regulation and spectral multiplexing. Nat Chem Biol 13(6):633–639.

4. Kennedy MJ, et al. (2010) Rapid blue-light-mediated induction of protein interactions in living cells. Nat Methods 7(12):973–975.

5. Toettcher JE, Gong D, Lim WA, & Weiner OD (2011) Light control of plasma membrane recruitment using the Phy-PIF system. Methods Enzymol 497:409–423.

6. Dagliyan O, et al. (2016) Engineering extrinsic disorder to control protein activity in living cells. Science 354(6318):1441–1444.

7. Guntas G, et al. (2015) Engineering an improved light-induced dimer (iLID) for controlling the localization and activity of signaling proteins. Proc Natl Acad Sci U S A 112(1):112–117.

8. Quejada JR, et al. (2017) Optimized light-inducible transcription in mammalian cells using Flavin Kelch-repeat F-box1/GIGANTEA and CRY2/CIB1. Nucleic Acids Res 45(20):e172.

9. Oliinyk OS, Shemetov AA, Pletnev S, Shcherbakova DM, & Verkhusha VV (2019) Smallest near-infrared fluorescent protein evolved from cyanobacteriochrome as versatile tag for spectral multiplexing. Nat Commun 10(1):279.

10. Reis JM, et al. (2018) Discovering Selective Binders for Photoswitchable Proteins Using Phage Display. ACS Synth Biol 7(10):2355–2364.

11. Wang H, et al. (2016) LOVTRAP: an optogenetic system for photoinduced protein dissociation. Nat Methods 13(9):755–758.

12. Kolar K, Knobloch C, Stork H, Znidaric M, & Weber W (2018) OptoBase: A Web Platform for Molecular Optogenetics. ACS Synth Biol 7(7):1825–1828.

13. Ikeuchi M & Ishizuka T (2008) Cyanobacteriochromes: a new superfamily of tetrapyrrole-binding photoreceptors in cyanobacteria. Photochem Photobiol Sci 7(10):1159–1167.

14. Fushimi K & Narikawa R (2019) Cyanobacteriochromes: photoreceptors covering the entire UV-to-visible spectrum. Curr Opin Struct Biol 57:39–46.

15. Anders K & Essen LO (2015) The family of phytochrome-like photoreceptors: diverse, complex and multi-colored, but very useful. Curr Opin Struct Biol 35:7–16.

16. Rockwell NC, Martin SS, Feoktistova K, & Lagarias JC (2011) Diverse two-cysteine photocycles in phytochromes and cyanobacteriochromes. Proc Natl Acad Sci U S A 108(29):11854–11859.

17. Rockwell NC, Martin SS, Gan F, Bryant DA, & Lagarias JC (2015) NpR3784 is the prototype for a distinctive group of red/green cyanobacteriochromes using alternative Phe residues for photoproduct tuning. Photochem Photobiol Sci 14(2):258–269.

18. Rockwell NC, Martin SS, & Lagarias JC (2012) Red/green cyanobacteriochromes: sensors of color and power. Biochemistry 51(48):9667–9677.

19. Rockwell NC, Martin SS, & Lagarias JC (2016) Identification of Cyanobacteriochromes Detecting Far-Red Light. Biochemistry 55(28):3907–3919.

20. Fushimi K, Ikeuchi M, & Narikawa R (2017) The Expanded Red/Green Cyanobacteriochrome Lineage: An Evolutionary Hot Spot. Photochem Photobiol 93(3):903–906.

21. Narikawa R, Enomoto G, Ni Ni W, Fushimi K, & Ikeuchi M (2014) A new type of dual-Cys cyanobacteriochrome GAF domain found in cyanobacterium Acaryochloris marina, which has an unusual red/blue reversible photoconversion cycle. Biochemistry 53(31):5051–5059.

22. Cornilescu CC, et al. (2014) Dynamic structural changes underpin photoconversion of a blue/green cyanobacteriochrome between its dark and photoactivated states. J Biol Chem 289(5):3055–3065.

23. Lim S, et al. (2014) Photoconversion changes bilin chromophore conjugation and protein secondary structure in the violet/orange cyanobacteriochrome NpF2164g3’ [corrected]. Photochem Photobiol Sci 13(6):951–962.

24. Narikawa R, et al. (2013) Structures of cyanobacteriochromes from phototaxis regulators AnPixJ and TePixJ reveal general and specific photoconversion mechanism. Proc Natl Acad Sci U S A 110(3):918–923.

25. Lim S, et al. (2018) Correlating structural and photochemical heterogeneity in cyanobacteriochrome NpR6012g4. Proc Natl Acad Sci U S A 115(17):4387–4392.

26. Blain-Hartung M, et al. (2018) Cyanobacteriochrome-based photoswitchable adenylyl cyclases (cPACs) for broad spectrum light regulation of cAMP levels in cells. J Biol Chem 293(22):8473–8483.

27. Fushimi K, et al. (2016) Photoconversion and Fluorescence Properties of a Red/Green-Type Cyanobacteriochrome AM1_C0023g2 That Binds Not Only Phycocyanobilin But Also Biliverdin. Front Microbiol 7:588.

28. Kainrath S, Stadler M, Reichhart E, Distel M, & Janovjak H (2017) Green-Light-Induced Inactivation of Receptor Signaling Using Cobalamin-Binding Domains. Angew Chem Int Ed Engl 56(16):4608–4611.

29. Chatelle C, et al. (2018) A Green-Light-Responsive System for the Control of Transgene Expression in Mammalian and Plant Cells. ACS Synth Biol 7(5):1349–1358.

30. Ong NT & Tabor JJ (2018) A Miniaturized Escherichia coli Green Light Sensor with High Dynamic Range. Chembiochem 19(12):1255–1258.

31. Chernov KG, Redchuk TA, Omelina ES, & Verkhusha VV (2017) Near-Infrared Fluorescent Proteins, Biosensors, and Optogenetic Tools Engineered from Phytochromes. Chem Rev 117(9):6423–6446.

32. Shu X, et al. (2009) Mammalian expression of infrared fluorescent proteins engineered from a bacterial phytochrome. Science 324(5928):804–807.

33. Fellouse FA & Sidhu SS (2007) Making antibodies in bacteria. Making and using antibodies: A practical handbook, eds Howard GC & Kaser MR (CRC Press, Boca Raton, FL), pp 157–180.

34. Keller S, et al. (2012) High-precision isothermal titration calorimetry with automated peak-shape analysis. Anal Chem 84(11):5066–5073.

35. Brautigam CA, Zhao H, Vargas C, Keller S, & Schuck P (2016) Integration and global analysis of isothermal titration calorimetry data for studying macromolecular interactions. Nat Protoc 11(5):882–894.

36. Zhao H, Piszczek G, & Schuck P (2015) SEDPHAT--a platform for global ITC analysis and global multi-method analysis of molecular interactions. Methods 76:137–148.

37. Kelley LA, Mezulis S, Yates CM, Wass MN, & Sternberg MJ (2015) The Phyre2 web portal for protein modeling, prediction and analysis. Nat Protoc 10(6):845–858.

38. Gambetta GA & Lagarias JC (2001) Genetic engineering of phytochrome biosynthesis in bacteria. Proc Natl Acad Sci U S A 98(19):10566–10571.

39. Song C, et al. (2015) A Red/Green Cyanobacteriochrome Sustains Its Color Despite a Change in the Bilin Chromophore’s Protonation State. Biochemistry 54(38):5839–5848.

40. Xu XL, et al. (2014) Combined mutagenesis and kinetics characterization of the bilin-binding GAF domain of the protein Slr1393 from the Cyanobacterium Synechocystis PCC6803. Chembiochem 15(8):1190–1199.

41. Goglia AG, Wilson MZ, DiGiorno DB, & Toettcher JE (2017) Optogenetic Control of Ras/Erk Signaling Using the Phy-PIF System. Methods Mol Biol 1636:3–20.

42. Uda Y, et al. (2017) Efficient synthesis of phycocyanobilin in mammalian cells for optogenetic control of cell signaling. Proc Natl Acad Sci U S A 114(45):11962–11967.

43. Fairbrother WJ, et al. (1998) Novel peptides selected to bind vascular endothelial growth factor target the receptor-binding site. Biochemistry 37(51):17754–17764.

44. Kirpich JS, et al. (2019) Reverse Photodynamics of the Noncanonical Red/Green NpR3784 Cyanobacteriochrome from Nostoc punctiforme. Biochemistry 58(18):2307–2317.

45. Pathak GP, Strickland D, Vrana JD, & Tucker CL (2014) Benchmarking of optical dimerizer systems. ACS Synth Biol 3(11):832–838.

46. Li L & Lagarias JC (1994) Phytochrome assembly in living cells of the yeast Saccharomyces cerevisiae. Proc Natl Acad Sci U S A 91(26):12535–12539.

47. Fushimi K, et al. (2019) Rational conversion of chromophore selectivity of cyanobacteriochromes to accept mammalian intrinsic biliverdin. Proc Natl Acad Sci U S A 116(17):8301–8309.

