## Supplemental Info for "Green, orange, red, and far-red optogenetic tools derived from cyanobacteriochromes"

### Supplementary Information

**Supplementary Methods:**

*Expression of Am1_c0023g2 and in vitro reconstitution*

For production of *Am1_c0023g2* for *in vitro* characterization, the gene was subcloned into the pET24b vector containing a C-terminal poly His (6x) tag. Transformed BL21(DE3) *E. coli* were grown at 37°C in LB medium (+50 mg/mL kanamycin) until OD 0.6, and expression was induced with 1 mM of IPTG. Cells were then grown at 18°C with shaking for 16 h. Cells were harvested, then sonicated in lysis buffer (50 mM phosphate, 300 mM NaCl, pH 7.5). The supernatant was separated by centrifugation, and applied to a Ni-NTA column. This was washed with lysis buffer + 15 mM imidazole, and subsequently eluted with lysis buffer + 100 mM EDTA. Eluted apo *Am1_c0023g2* was dialyzed against PBS (137 mM NaCl, 3 mM KCl, 8 mM $Na_2HPO_4$, 1.5 mM $KH_2PO_4$, pH 7.2 (+ 5 mM β-ME). β-ME was then removed using an Amicon 10K spin filter, after which the apo protein was loaded back onto an Ni-NTA column. A two-fold molar excess of purified phycocyanobilin (PCB, Frontier Scientific) or biliverdin (BV, Frontier Scientific) in DMSO was added to the Ni-NTA column equilibrated in lysis buffer containing the purified apo protein and incubated overnight at 4°C. Holo-protein was eluted using lysis buffer + 100 mM EDTA, then further purified using a Superdex 75 10/300 GL size exclusion column (GE Healthcare) (running in PBS at 0.4 mL/min)

For phage display, *Am1_c0023g2* was cloned into an expression vector containing an N-terminal GST-Avi-TEV site with a C-terminal 6X His-tag (1). AVB100 *E. Coli* K12 cells (AVIDITY) were used for *in vivo* biotinylation during recombinant expression of the protein. Cells were grown at 37°C in LB medium (+100 µg/mL ampicillin) until OD 0.6, which was then supplemented with 50 µM D-biotin, 0.2% L-arabinose, and 1 mM IPTG. Cells were grown at 18°C for 14 h. Protein purification was carried out as described above.

*Expression of Am1_c0023g2 PCB by in vivo reconstitution*

For the expression of *Am1_c0023g2* PCB with *in vivo* chromophore reconstitution, the gene was subcloned into a pBAD-HisC vector with C-terminal 6x His-tag. *E. coli* strain LMG194 containing pPL-PCB (a kind gift from J.C. Lagarias) was transformed with pBAD-*Am1_c0023g2* and cells exhibiting both kanamycin and ampicillin resistance were selected. Cells were grown overnight in 1 mL RM media (2% casamino acids (BD Biosciences, without iron), 1X M9 salts, 1 mM $MgCl_2$) with 0.2% glucose, 50 µg/mL kanamycin, and 100 µg/mL ampicillin at 37$^o$C. A 1 mL volume of the overnight culture was added to 100 mL of RM media containing 0.2% glucose, 50 µg/mL kanamycin, and 100 µg/mL ampicillin and incubated at 37$^o$C (180 rpm). When OD 0.5 was reached, the 100 mL culture was added to 900 mL LB media (+ 50 mg kanamycin, 100 mg ampicillin, 1 mM IPTG) and grown for 1 hour at 37$^o$C (180 rpm). At that point, 0.002% L-arabinose (20 mg) was added and the culture was incubated at 37$^o$C for a further 4 h (180 rpm), then cells were harvested by centrifugation. Cells were sonicated in lysis buffer (50 mM phosphate, 300 mM NaCl, pH 7.2), centrifuged to remove cell debris, and the supernatant was applied to a Ni-NTA column. The column was washed with lysis buffer containing 15 mM imidazole and the protein was eluted with lysis buffer containing 100 mM EDTA. Eluted protein was dialyzed against 1x PBS (137 mM NaCl, 3 mM KCl, 8 mM $Na_2HPO_4$, 1.5 mM $KH_2PO_4$, pH 7.2) and purified further using a Superdex 75 10/300 GL size exclusion column (GE Healthcare) in PBS (0.4 mL/min flow rate).

*Expression of Am1_c0023g2 BV by in vivo reconstitution*

For the expression of *Am1_c0023g2* BV with *in vivo* chromophore reconstitution, the gene was subcloned into a pBAD-HisC vector with a C-terminal 6x His-tag. *E. coli* strain LMG194 containing pWA23h (a kind gift from Prof. Vladislav Verkhusha) was transformed with pBAD-*Am1_c0023g2* and cells exhibiting both kanamycin and ampicillin resistance were selected. Cells were grown overnight in 10 mL RM media (2% casamino acids (BD Biosciences), 1X M9 salts, 1 mM $MgCl_2$) with 0.2% glucose, 50 µg/mL kanamycin, and 100 µg/mL ampicillin at 37ºC. A 5 mL volume of the overnight culture was added to 1 L of RM media containing 0.02% L-rhamnose, 50 µg/mL kanamycin, and 100 µg/mL ampicillin and incubated at 37ºC (180 rpm) until OD 0.6 was reached. Expression of *Am1_c0023g2* was then induced with 0.002% L-arabinose then cultures were incubated at 30ºC for 12 h (180 rpm), followed by incubation at 18 ºC for 12 h (180 rpm). Purification of *Am1_c0023g2* BV followed the same protocol described above for purification of Am1_c0023g2 PCB

*Phage display-based screening*

A M13 phage pVIII library based on the GA domain described previously (2) was used to find binders for the Pg or Pr state of *Am1_c0023g2* PCB and the Po or Pfr state of *Am1_c0023g2* BV using the following protocol. MaxiSorp 96 well plates were coated with 5 µg/mL neutravidin (in PBS) and blocked with PB buffer (PBS, 2 mg/mL BSA). 20 µg/mL of biotinylated apo- or holo-*Am1_c0023g2* (in PBS + 2 mg/mL BSA + 0.05% Tween-20) was added and incubated for 2 h at room temperature. After the removal of any unbound protein, plates were either placed under green (525 nm, 1 mW/cm$^2$) to produce the Pr state, red (680 nm, 2.8 mW/cm$^2$) to produce the Pg state, far-red (750 nm to produce the Po state (1.1 mW/cm$^2$), or dark (to produce the Pfr state) for 1 hour. Apo plates were placed under ambient light during this step. The GA domain phage library (~5 x $10^{12}$ cfu/mL) was added to the apo plate first. After incubation for 1 hour at room temperature, the supernatant containing unbound phage was transferred to either a Pg, Pr, Po or Pfr state plate for a second negative selection, followed by a positive selection on a Pr, Pg, Pfr or Po state plate, respectively, for 1 hour each at room temperature. Following an extensive wash (8 times), with PBS + 0.02% Tween-20, the bound phage was eluted and amplified following standard protocols (3). The whole process was repeated two times including apo and negative selections. The resulting library of positive clones obtained after 3 rounds of selection was subcloned into a pIII phagemid using *SacI* and *NsiI* as described previously (2). At this point ~100 single clonal phage from each of the positive pools were screened for binding to either the Pg or the Pr state of *Am1_c0023g2* PCB and Pfr or Po state of *Am1_c0023g2* BV via a phage-based ELISA (3).

*Affinity Maturation*

Biased libraries were constructed in the pIII display format, based on doped oligonucleotides specific for lead clones BAm-green 1.0 and BAm-red 1.0. Single-stranded DNA was isolated from each clone and used as a template for site directed mutagenesis (4). We used the following oligonucleotides for mutagenesis.

BAm-green 1.0

5’-AAGGCTGGTATCACC(N3)(N1)(N4)GAC(N2)(N2)(N4)(N2)(N3)(N4)TTCAAC(N3)(N2)(N4)ATCAAT(N4)(N4)(N4)GCG(N4)(N4)(N3)(N3)(N1)(N4)GTG(N1)(N1)(N4)(N4)(N4)(N4)GTTAAC(N1)(N1)(N3)(N4)(N4)(N4)AAGAAC(N4)(N1)(N4)ATCCTGAAAGCTCAC-3’

BAm-red 1.0

5’-AAGGCTGGTATCACC(N1)(N1)(N4)GAC(N4)(N3)(N3)(N4)(N4)(N4)TTCAAC(N3)(N1)(N4)ATCAAT(N4)(N2)(N4)GCG(N4)(N4)(N4)(N4)(N1)(N4)GTG(N4)(N2)(N4)(N3)(N1)(N4)GTTAAC(N3)(N4)(N4)(N4)(N4)(N3)AAGAAC(N4)(N1)(N4)ATCCTGAAAGCTCAC-3’

Where N1 is a mix of 70% A, 10% C, 10% G, 10% T
N2 is a mix of 10% A, 70% C, 10% G, 10% T
N3 is a mix of 10% A, 10% C, 70% G, 10% T
N4 is a mix of 10% A, 10% C, 10% G, 70% T

Libraries were constructed using previously published protocols and resulted in a diversity of approximately $10^9$ different clones (5). Affinity maturation selection was performed using the same protocol as naïve selection except streptavidin was used at 2 µg/mL and 5 µg/mL of *Am1_c0023g2* PCB was added to each well.

*Expression of binders:*

Selected binders were subcloned from pIII phagemids into the pET24b expression vector containing a C-terminal poly His (6x) tag and transformed into BL21 (DE3) cells. Cells were grown until mid-log phase (OD ~0.6), and then induced via addition of 0.75 mM IPTG and grown at 20$^{o}$C for 18 h (180 rpm). Cells were harvested, sonicated in lysis buffer (50 mM phosphate, 300 mM NaCl, pH 7.2), and centrifuged to remove cell debris. The supernatant was loaded onto a Ni-NTA column, washed with lysis buffer containing 15 mM imidazole, and eluted using lysis buffer containing 250 mM imidazole. Following elution from the Ni-NTA column, proteins were dialyzed against 1X PBS (pH 7.2) and further purified using a Superdex 75 10/300 GL size exclusion column (GE Healthcare) in PBS (137 mM NaCl, 3 mM KCl, 8 mM $Na_2HPO_4$, 1.5 mM $KH_2PO_4$, pH 7.2, 0.4 mL/min flow rate)

*Size exclusion binding assay*

*Am1_c0023g2* PCB or BV (80 µM) was mixed with each binder (80 µM) in PBS (137 mM NaCl, 3 mM KCl, 8 mM $Na_2HPO_4$, 1.5 mM $KH_2PO_4$, pH 7.2) and injected onto a Superdex 75 10/300 GL size exclusion column (GE Healthcare, 0.4 mL/min flow rate) maintained in the dark or while irradiating with either red (680 nm), green (525 nm), or far red (750 nm) light both to the column and sample. Eluted proteins were collected in 0.45 mL fractions, each of which was electrophoresed on a 12.5% SDS-PAGE gel. The column was calibrated using size standards (3 mg/ml each of conalbumin, carbonic anhydrase, ribonuclease A, and aprotinin).

*UV-Vis spectroscopy measurements*

UV-Vis spectra were obtained using Perkin Elmer Lambda 35 spectrophotometer with a temperature-controlled cuvette holder (Quantum Northwest). To a sample *Am1_c0023g2* PCB or BV (10 µM final concentration) irradiated with 680 nm or 750 nm, binders were added in the concentrations indicated. Absorbance spectra were acquired at 1 hour intervals for 24h at $20^{o}$C.

*Fluorescence quenching binding measurements*

To a fixed amount of *Am1_c0023g2* PCB exo (3 µM) in 1X PBS (pH 7.2), increasing amounts of BAm-red1.0 or *wt*-GA domain (1 to 20 µM) were added, and the samples were irradiated with a 525 nm LED (to produce the Pr state) for 30 minutes. Samples were excited at 628 nm and fluorescence emission was collected at 678 ± 37 nm using a BMG Labtech Clariostar plate reader. Fluorescence intensity data was fit to a Morrison equation for tight binding:

$$\mathrm{I}(B_{tot}) = I_{init} + \left(I_{final} - I_{init}\right)\left[\frac{(B_{tot} + A_{tot} + K_d) - \sqrt{(-B_{tot} - A_{tot} - K_d)^2 - (4 * A_{tot} * B_{tot})}}{2 * A_{tot}}\right]$$

where $A_{tot} = [Am1\_c0023g2]$ $B_{tot} = [BAm - red1.0]$

*Isothermal titration calorimetry (ITC)*

ITC experiments were performed using a MicroCal VP-ITC MicroCalorimeter. Samples prepared as described above for size exclusion chromatography binding assays. Titration experiments were performed at 25°C. The syringe contained 250 µL of the binder, BAm-red1.0 or BAm-green1.3, at 550 µM or 750 µM, respectively. The cell (1.4 mL) contained either *Am1_c0023g2* PCB exo (45 µM), *Am1_c0023g2* PCB endo (42 µM), *Am1_c0023g2* BV exo (45 µM) or *Am1_c0023g2* BV endo (42 µM). To obtain the Pg state (PCB loaded protein) samples were pre irradiated at 680 nm. To obtain the Pr state, samples were pre-irradiated at 525 nm prior to transferring into the cell. For *Am1_c0023g2* BV, samples were incubated in dark for 3 h to adapt to the Pfr state. A small amount of the undesired state was formed in each case due to a brief exposure to low intensity white light required for loading the syringe into the instrument. All samples were equilibrated with 1X PBS, pH 7.2. Injections volumes were 10 µL for BAm-red1.0 and 5 µL for BAm-green1.3 with a 300 s spacing between injections. Thermogram data were integrated using NITPIC (6) and binding analysis carried out using SEDPHAT using recommend protocols (7, 8).

*Construction of yeast plasmids*

The strains Y2HGOLD (*MATa, trp1-901, leu2-3, 112, ura3-52, his3-200, gal4Δ, gal80Δ, LYS2 : :GAL1*$_{UAS}$*-GAL1*$_{TATA}$*-His3, GAL2*$_{UAS}$*-Gal2*$_{TATA}$*-Ade2, URA3 : : MEL1*$_{UAS}$*-Mel1*$_{TATA}$*, AUR1-C MEL1*) and Y187 (*MATα, ura3-52, his3-200, ade2-101, trp-901, leu2-3, 112, gal4Δ, gal80Δ, met-, URA3 : : GAL1*$_{UAS}$*-Gal1*$_{TATA}$*-LacZ, MEL1*) were purchased from Clontech. pGAL4AD-x and pGAL4BD-y plasmids were purchased from Addgene (28246 and 28244) as pGAL4AD-CIB1 and pGAL4BD-Cry2 (9). The *Am1_c0023g2* gene was PCR amplified and inserted (Gibson Assembly) into either pGAL4AD or pGAL4BD vector by replacing a CIB1 or Cry2 gene, respectively. Binder constructs were also subcloned into both plasmids using the same protocol.

*β-galactosidase assay*

Y187 and Y2HGOLD strains containing pGAL4AD-x and pGAL4BD-y, respectively, were mated in 5 mL of YPDA (1% yeast extract, 2% peptone, 2% glucose, 0.02% adenine) overnight. Mated clones were selected on SC -L/W (synthetic media lacking leucine and tryptophan). SC -L/W media contained 2 g/L drop out medium (Bioshop; no uracil, histidine, leucine, tryptophan, adenine), 2% glucose, 50 mg/L histidine, 100 mg/L adenine hemisulfate, 20 mg/L uracil, 1.7 g/L yeast nitrogen base (Biobasic), 5 g/L ammonium sulfate (Bioshop). A single colony was picked and used to inoculate 5 mL of SC -L/W then grown at 30$^{o}$C for 36 h (160 rpm). Following growth, cells were diluted to $OD_{600nm}$ = 0.2 in 1.2 mL SC -L/W media containing 10 µM PCB (Frontier Scientific). Cultures were grown in the dark for 3 h, then grown with either red (680 nm, 81 µW/cm$^2$) or green (525 nm, 58 µW/cm$^2$) irradiation for 4 more hours These intensities did not appear to cause any significant bleaching of PCB in solution over this timeframe. After the 7-hour growth period, 800 µL of cells were harvested and washed one time with Z-buffer (Clontech Laboratories), followed by lysis with Y Cell Lytic reagent (Sigma Aldrich). β-galactosidase activity was measured following the protocol from Matchmaker Gold Yeast Two-Hybrid System (Takara, Clontech) using ONPG as a substrate. The experiment was performed in quadruplicate. Experiments with BV were performed in the same manner but with SC -L/W media containing 40 µM BV (Frontier Scientific). During the 4 h irradiation phase, cells were either incubated under 750 nm (1.1 $mW/cm^2$) or in the dark.

### Supplementary Figures:

```
NpR6012g4                 ------------------------------------------------------------ 0
Am1_c0023g2               ------------------------------------------------------------ 0
Shane_Am1_c0023g2_L405K   MKIEESSGKLMSPILGYWKIKGLVQPTRLLLEYLEEKYEEHLYERDEGDKWRNKKFELGL 60
Am1_c0023g2_L405K         ------------------------------------------------------------ 0

NpR6012g4                 ------------------------------------------------------------ 0
Am1_c0023g2               ------------------------------------------------------------ 0
Shane_Am1_c0023g2_L405K   EFPNLPYYIDGDVKLTQSMAIIRYIADKHNMLGGCPKERAEISMLEGAVLDIRYGVSRIA 120
Am1_c0023g2_L405K         ------------------------------------------------------------ 0

NpR6012g4                 ------------------------------------------------------------ 0
Am1_c0023g2               ------------------------------------------------------------ 0
Shane_Am1_c0023g2_L405K   YSKDFETLKVDFLSKLPEMLKMFEDRLCHKTYLNGDHVTHPDFMLYDALDVVLYMDPMCL 180
Am1_c0023g2_L405K         ------------------------------------------------------------ 0

NpR6012g4                 ------------------------------------------------------------ 0
Am1_c0023g2               --------------------------------------------------MGSSHHHHHH 10
Shane_Am1_c0023g2_L405K   DAFPKLVCFKKRIEAIPQIDKYLKSSKYIAWPLQGWQATFGGGDHPPKSDLEVLFQGPLS 240
Am1_c0023g2_L405K         ------------------------------------------------------------ 0

NpR6012g4                 --------------MGEKAVTKISNRIRQSSDVEEIFKTTTQEVRQLLRCDRVAVYRFNP 46 (628)
Am1_c0023g2               S-GSGLVP----RGSHMRNISEIIQRIRQSLDIEDIFGATTQDVRESLECDRVVIYQFWP 65 (279)
Shane_Am1_c0023g2_L405K   SGGGGLNDIFEAQKIEWHEEDLYFQSAAKSLDIEDIFGATTQDVRESLECDRVVIYQFWP 300
Am1_c0023g2_L405K         ---------------------------MKSLDIEDIFGATTQDVRESLECDRVVIYQFWP 33 (279)
                                                    :* *:*:** :***:**: *.****.:*:* *
NpR6012g4                 NWTGEFVAESVAHTWVKLVGPDIKTVWEDTHLQETQGGRYAQGENFVVNDIYQVGHSPCH 106 (688)
Am1_c0023g2               DWSGEFLVESTAPGLIPLSELDVPMTWQDTYLQENQGGKFKDNAPTVVADIYQQSYTDCH 125 (339)
Shane_Am1_c0023g2_L405K   DWSGEFLVESTAPGLIPLSELDVPMTWQDTYLQENQGGKFKDNAPTVVADIYQQGYTDCH 360
Am1_c0023g2_L405K         DWSGEFLVESTAPGLIPLSELDVPMTWQDTYLQENQGGKFKDNAPTVVADIYQQGYTDCH 93 (339)
                          :*:***:.**.*   : *   *:  .*:**:***.***:: :.   ** ****  .:: **
NpR6012g4                 IEILEQFEVKAYVIVPVFAGEQLWGLLAAYQNSGTRDWDESEVTLLARIGNQLGLALQQT 166 (748)
Am1_c0023g2               LEILEWFDIRAYMVVPVFIGKTLWGLLAAYQLNHPRQWQKVELYLLKQAGAQLGVALQQA 185 (399)
Shane_Am1_c0023g2_L405K   LEILEWFDIRAYMVVPVFIGKTLWGLLAAYQLNHPRQWQKVELYLLKQAGAQLGVALQQA 420
Am1_c0023g2_L405K         LEILEWFDIRAYMVVPVFIGKTLWGLLAAYQLNHPRQWQKVELYLLKQAGAQLGVALQQA 153 (399)
                          :**** *:::**::**** *: *********  .  *:*:: *: ** : * ***:****:
NpR6012g4                 EYLQQVQGQSAK--       178 (760)
Am1_c0023g2               ELLNQLR-------       192 (406)
Shane_Am1_c0023g2_L405K   ELLNQKLEHHHHHH       434
Am1_c0023g2_L405K         ELLNQKLEHHHHHH       167 (405)
                          * *:*
```

**GA domain sequences:**

| | | | | | | | | | | | | | | | | | | | | | | | | | | | | | | | | | | | | | | | | | | | | | | | | | | | | | |
|---|---|---|---|---|---|---|---|---|---|---|---|---|---|---|---|---|---|---|---|---|---|---|---|---|---|---|---|---|---|---|---|---|---|---|---|---|---|---|---|---|---|---|---|---|---|---|---|---|---|---|---|---|---|
| wt-GA-domain | T | I | D | Q | W | L | L | K | N | A | K | E | D | A | I | A | E | L | K | K | A | G | I | T | S | D | F | Y | F | N | A | I | N | K | A | K | T | V | E | E | V | N | A | L | K | N | E | I | L | K | A | H | A |
| Bam-red 1.0 | T | I | D | Q | W | L | L | K | N | A | K | E | D | A | I | A | E | L | K | K | A | G | I | T | N | D | W | F | F | N | D | I | N | S | A | F | Y | V | S | D | V | N | V | L | K | N | Y | I | L | K | A | H | A |
| Bam-red 1.1 | T | I | D | Q | W | L | L | K | N | A | K | E | D | A | I | A | E | L | K | K | A | G | I | T | K | D | W | Y | F | N | D | I | N | S | A | L | Y | V | S | D | V | N | V | L | K | N | H | I | L | K | A | H | A |
| Bam-red 1.2 | T | I | D | Q | W | L | L | K | N | A | K | E | D | A | I | A | E | L | K | K | A | G | I | T | N | D | W | F | F | N | D | I | N | S | A | L | F | V | S | D | V | N | V | L | K | N | H | I | L | K | A | H | A |
| Bam-red 1.3 | T | I | D | Q | W | L | L | K | N | A | K | E | D | A | I | A | E | L | K | K | A | G | I | T | L | D | W | Y | F | N | A | I | N | S | A | L | F | V | S | D | V | N | V | L | K | N | Y | I | L | K | A | H | A |
| Bam-red 1.4 | T | I | D | Q | W | L | L | K | N | A | K | E | D | A | I | A | E | L | K | K | A | G | I | T | I | D | W | Y | F | N | G | I | N | S | A | L | Y | V | T | D | V | N | V | L | K | N | Y | I | L | K | A | H | A |
| Bam-red 1.5 | T | I | D | Q | W | L | L | K | N | A | K | E | D | A | I | A | E | L | K | K | A | G | I | T | N | D | W | F | F | N | S | I | N | S | A | L | Y | V | T | D | V | N | A | L | K | N | H | I | L | K | A | H | A |
| Bam-green 1.0 | T | I | D | Q | W | L | L | K | N | A | K | E | D | A | I | A | E | L | K | K | A | G | I | T | D | D | P | R | F | N | A | I | N | F | A | L | D | V | N | F | V | N | K | F | K | N | Y | I | L | K | A | H | A |
| Bam-green 1.1 | T | I | D | Q | W | L | L | K | N | A | K | E | D | A | I | A | E | L | K | K | A | G | I | T | C | D | P | R | F | N | A | I | N | Y | A | R | D | V | D | F | V | N | K | F | K | N | Y | I | L | K | A | H | A |
| Bam-green 1.2 | T | I | D | Q | W | L | L | K | N | A | K | E | D | A | I | A | E | L | K | K | A | G | I | T | D | D | P | R | F | N | A | I | N | Y | A | K | D | V | N | F | V | N | K | F | K | N | Y | I | L | K | A | H | A |
| Bam-green 1.3 | T | I | D | Q | W | L | L | K | N | A | K | E | D | A | I | A | E | L | K | K | A | G | I | T | D | D | P | R | F | N | A | I | N | Y | A | L | S | V | E | F | V | N | K | F | K | N | Y | I | L | K | A | H | A |
| Bam-green 1.4 | T | I | D | Q | W | L | L | K | N | A | K | E | D | A | I | A | E | L | K | K | A | G | I | T | C | D | P | R | F | N | A | I | N | F | A | R | D | V | N | F | V | N | K | F | K | N | H | I | L | K | A | H | A |
| Bam-green 1.5 | T | I | D | Q | W | L | L | K | N | A | K | E | D | A | I | A | E | L | K | K | A | G | I | T | D | D | P | R | F | N | A | I | N | T | A | L | D | V | N | F | V | N | K | F | K | N | Y | I | L | K | A | H | A |
| BV clones | | | | | | | | | | | | | | | | | | | | | | | | | | | | | | | | | | | | | | | | | | | | | | | | | | | | | |
| D7 | T | I | D | Q | W | L | L | K | N | A | K | E | D | A | I | A | E | L | K | K | A | G | I | T | S | D | A | Y | F | N | F | I | N | D | A | L | V | V | S | S | V | N | V | R | K | N | Y | I | L | K | A | H | A |
| B5 | T | I | D | Q | W | L | L | K | N | A | K | E | D | A | I | A | E | L | K | K | A | G | I | T | D | D | L | T | F | N | V | I | N | N | A | Y | S | V | F | M | V | N | T | K | K | N | S | I | L | K | A | H | A |
| C3 | T | I | D | Q | W | L | L | K | N | A | K | E | D | A | I | A | E | L | K | K | A | G | I | T | D | D | A | V | F | N | F | I | N | S | A | Y | L | V | V | S | V | N | T | R | K | N | H | I | L | K | A | H | A |
| E6 | T | I | D | Q | W | L | L | K | N | A | K | E | D | A | I | A | E | L | K | K | A | G | I | T | S | D | F | H | F | N | S | I | N | S | A | L | F | V | S | D | V | N | L | V | K | N | S | I | L | K | A | H | A |

The expressed sequences had additions at the C- and N-terminal ends as follows:

wt> MKLATIDQWLLKNAKEDAIAELKKAGITSDFYFNAINKAKTVEEVNALKNEILKAHAGSSGLEHHHHHH

**Number of amino acids:** 69

**Molecular weight:** 7727.78

**Theoretical pI:** 7.07

**Figure S1. (top)** Sequence alignments of various constructs of *Am1_c0023g2*. *Shane_Am1_c0023g2_L405K* is a construct containing GST and Avi tag used for phage display. *Am1_c0023g2** is the version of the protein in Fushimi *et al.* (10). *Am1_c0023g2_L405K* is the engineered version used in the current work. Amino acids in red indicate the GAF domain region in each construct and the numbering in the full protein. The L405K mutation is highlighted in green. The expressed apoprotein used in this work has 167 amino acids, MW 19320.92, and a pI of 4.89. **(bottom)** GA domain sequences selected using phage display. When not displayed on phage, the proteins have short additions at the N- and C-terminal ends to permit *E. coli* expression and purification.

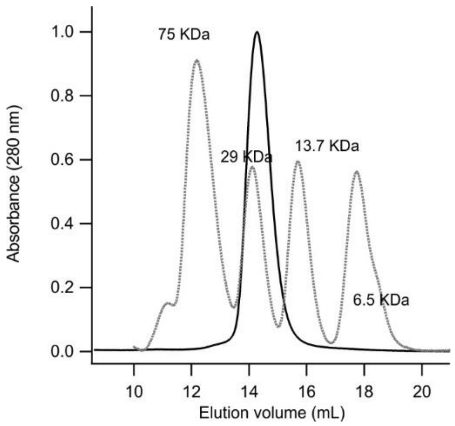

**Figure S2** Size exclusion chromatography of *Am1_c0023g2* PCB (solid line) was performed using a Superdex 75 10/300 GL column (GE Healthcare) on a BioRad Duoflow F10 workstation (flow rate 0.5 mL/min). The running buffer used was 137 mM NaCl, 3 mM KCl, 8 mM $Na_2HPO_4$, 1.5 mM $KH_2PO_4$, pH 7.2. The protein concentration was 75 µM. A mixture of 3 mg/mL conalbumin, carbonic anhydrase, ribonuclease A and aprotinin was used as a molecular weight calibration standard (dashed lines). The apparent molecular weight of *Am1_c0023g2* PCB ~22 kDa corresponds to a monomer.

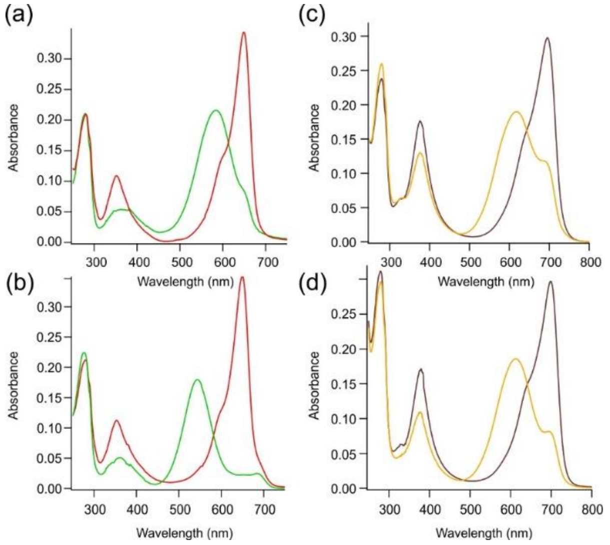

**Figure S3** (a) Absorbance spectra of *Am1_c0023g2* with exogenous PCB incorporation in the Pr and Pg states. (b) Absorbance spectra of *Am1_c0023g2* with endogenous PCB incorporation in the Pr and Pg states. (c) Absorbance spectra of *Am1_c0023g2* with exogenous BV incorporation in the Pfr and Po states. (d) Absorbance spectra of *Am1_c0023g2* with endogenous BV incorporation in the Pfr and Po states.

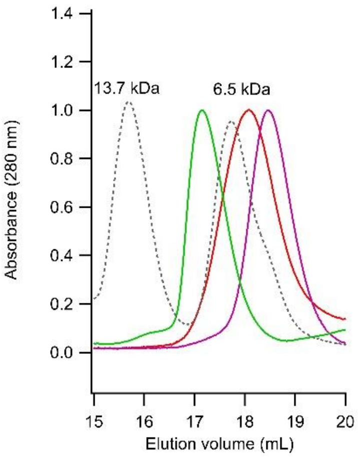

**Figure S4**. Size exclusion chromatography of binders was performed using a Superdex 75 10/300 GL column (GE Healthcare) on a BioRad Duoflow F10 workstation (flow rate 0.5 mL/min). The running buffer used was 137 mM NaCl, 3 mM KCl, 8 mM $Na_2HPO_4$, 1.5 mM $KH_2PO_4$, pH 7.2. The protein concentration was ~50 μM.  A mixture of 3 mg/mL conalbumin, carbonic anhydrase, ribonuclease A and aprotinin was used as a molecular weight calibration standard (dashed lines). The apparent molecular weights of the binders (all near 7 kDa) correspond to monomers.

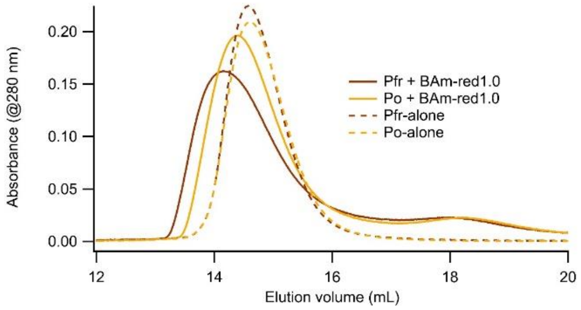

**Figure S5**. BV-loaded *Am1_c0023g2* shows complex formation with BAm-red1.0 in the Pfr state. Residual Pfr state binding is also observed (when elution is monitored at 280 nm) from the Po state sample (compare to Fig. 3c, main text).

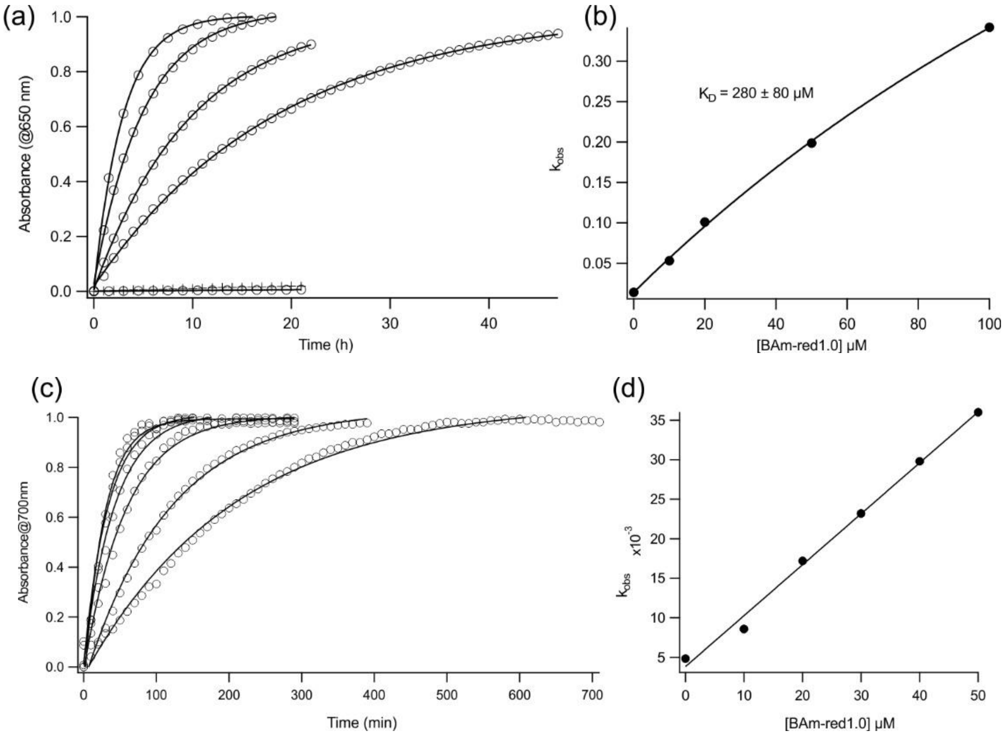

**Figure S6.** (a) UV-Vis measurement of the thermal relaxation of *Am1_c0023g2* PCB exo (10 µm) in the presence of increasing concentrations of BAm-red1.0 (0, 10, 20, 50, 100 µM) (circles). Wild-type GA domain (+) (50 µM) has no effect on the relaxation. (b) Rate constants obtained by fitting the curves in (a) to single exponential decay processes are plotted versus the concentration of added BAm-red1.0. When these data are fit to a Morrison Eq. for tight binding, a $K_D$ of 280 µM is obtained. (c) UV-Vis measurement of the thermal relaxation of *Am1_c0023g2* BV exo (10 µm) in the presence of increasing concentrations of BAm-red1.0 (0, 10, 20, 50 µM) (circles). (d) Rate constants obtained by fitting the curves in (c) to single exponential decay processes are plotted versus the concentration of added BAm-red1.0. There is an approximately linear increase in $k_{obs}$ vs. [BAm-red1.0] indicating weak (>500 µM) binding.

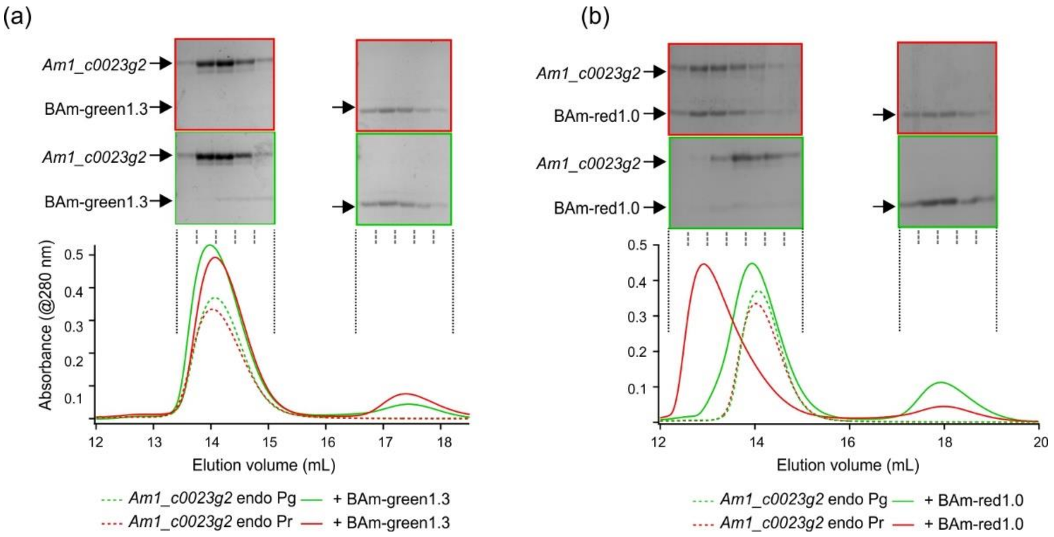

**Figure S7**. (a) PCB-loaded *Am1_c0023g2* endo shows no complex with BAm-green1.3 in the Pg state or the Pr state. (b) PCB-loaded *Am1_c0023g2* endo in the Pr state forms a complex with BAm-red1.0 and does not with the Pg state.

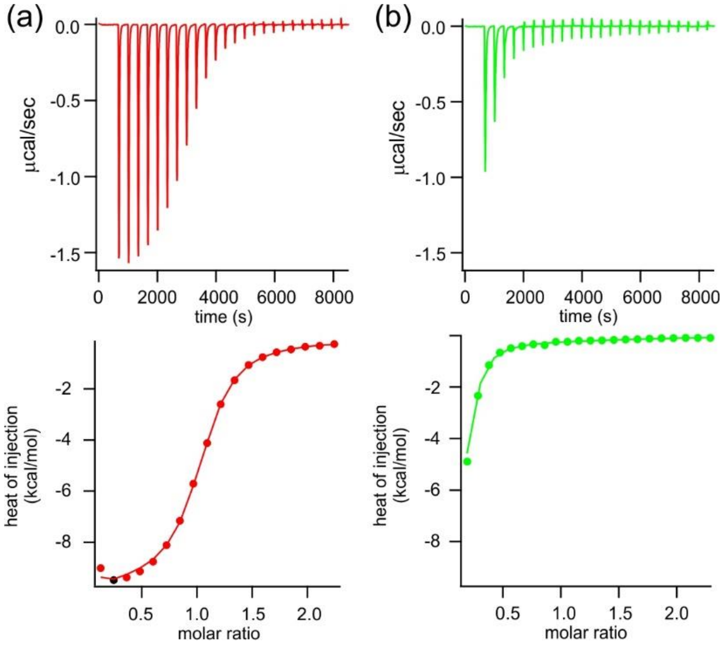

**Figure S8.** (a) Thermogram showing the titration of BAm-red1.0 (550 µM in the syringe) into a solution of *Am1_c0023g2* PCB endo (34 µM) in the Pr state and thermogram data fitted to a 1:1 binding model to give $K_D$ = 0.8 µM (0.65 - 1 µM) and ΔH = -9 ± 0.3 kcal/mol. (b) BAm-red1.0 was titrated into a solution of *Am1_c0023g2* PCB endo (34 µM) in the Pg state and thermogram data fitted to a model in which 20% of the Pr state remains and the Pg state is inactive.

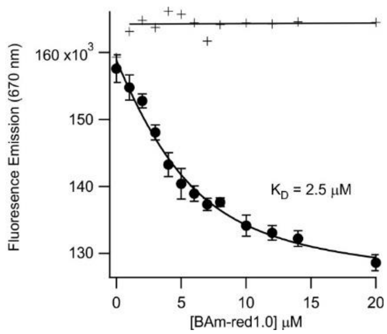

**Figure S9.** Fluorescence emission intensity (at 670 nm) of the Pr state of *Am1_c0023g2* PCB exo as a fucntion of added BAm-red1.0 (solid circles) or wild-type GA domain (+). Fitting the binding data to Eq. 1 gives a $K_D$ of 2.5 µM.

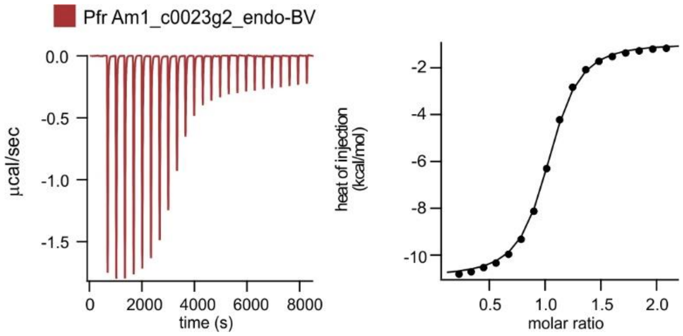

**Figure S10.** (left) Thermogram showing the titration of BAm-red1.0 (550 µM in the syringe) into a solution of *Am1_c0023g2* BV endo (36 µM) in the Pfr state. (right) The thermogram data were fitted to a 1:1 binding model to give $K_D$ = 0.75 µM (0.71 – 0.84 µM) and $\Delta H$ = -10.2 ± 0.3 kcal/mol.

### References:

1. Hornsby M, *et al.* (2015) A High Through-put Platform for Recombinant Antibodies to Folded Proteins. *Mol Cell Proteomics* 14(10):2833-2847.
2. Reis JM, *et al.* (2018) Discovering Selective Binders for Photoswitchable Proteins Using Phage Display. *ACS Synth Biol* 7(10):2355-2364.
3. Fellouse FA & Sidhu SS (2007) Making antibodies in bacteria. *Making and using antibodies: A practical handbook*, eds Howard GC & Kaser MR (CRC Press, Boca Raton, FL), pp 157-180.
4. Kunkel TA (1985) Rapid and efficient site-specific mutagenesis without phenotypic selection. *Proc. Natl. Acad. Sci. U.S.A.* 82(2):488-492.
5. Chen G & Sidhu SS (2014) Design and generation of synthetic antibody libraries for phage display. *Methods Mol Biol* 1131:113-131.
6. Keller S, *et al.* (2012) High-precision isothermal titration calorimetry with automated peak-shape analysis. *Anal Chem* 84(11):5066-5073.

7. Brautigam CA, Zhao H, Vargas C, Keller S, & Schuck P (2016) Integration and global analysis of isothermal titration calorimetry data for studying macromolecular interactions. *Nat Protoc* 11(5):882-894.
8. Zhao H, Piszczek G, & Schuck P (2015) SEDPHAT--a platform for global ITC analysis and global multi-method analysis of molecular interactions. *Methods* 76:137-148.
9. Kennedy MJ, *et al.* (2010) Rapid blue-light-mediated induction of protein interactions in living cells. *Nat Methods* 7(12):973-975.
10. Fushimi K, *et al.* (2016) Photoconversion and Fluorescence Properties of a Red/Green-Type Cyanobacteriochrome AM1_C0023g2 That Binds Not Only Phycocyanobilin But Also Biliverdin. *Front Microbiol* 7:588.
